# Seasonal and diel environment effects on host-seeking behaviour in the European tick vector, *Ixodes ricinus*

**DOI:** 10.1101/2025.11.06.685884

**Authors:** Madeleine Noll, Richard Wall, Benjamin L. Makepeace, Hannah Rose Vineer

## Abstract

**Background:** Patterns of arthropod vector abundance and activity are key determinants of biting risk, and thus the risk of vector-borne pathogen transmission. Ticks are important vectors of animal and human pathogens worldwide. Their host-seeking activity (questing) directly influences the risk of exposure to ticks and their pathogens, but the drivers of this behaviour are poorly understood.

**Methods:** The concurrent seasonal and diel variation in *Ixodes ricinus* questing behaviour was investigated, for the first time, by blanket dragging every four hours over a 24-hour sampling cycle each month, from January to December 2022. Associations between tick density and environmental factors were evaluated by Generalised Linear Mixed Modelling (GLMM).

**Results:** A predominantly nocturnal questing activity pattern was observed. Activity was also moderated by seasonal moisture availability, being most evident in nymphs in months with high saturation deficit. The density of questing nymphs was positively associated with darkness, and negatively associated with saturation deficit.

**Conclusions:** The observed seasonal questing patterns are largely consistent with published data; however this is the first study to highlight the importance of nocturnal feeding and periods of high saturation deficit in the field. These commonly overlooked daily rhythms and environmental moderators of tick behaviour have important implications for assessing the relative hazard and potential exposure to ticks and their pathogens for different hosts, and for estimating abundance in scientific research and surveillance.

## 1 Background

Seasonal patterns of arthropod vector abundance and activity, and diel activity patterns (variation in activity over a 24-hour cycle) are key determinants of biting risk (interacting with host presence and behaviour), and thus the potential transmission of vector-borne pathogens (e.g. [1]). Ticks are some of the most important vectors world-wide, transmitting a range of protozoal, bacterial and viral pathogens as they feed on the blood of their hosts. *Ixodes ricinus*, also known as the castor bean or sheep tick, is the most widespread and abundant tick species in the Western Palearctic [2]. The temporal hazard of tick-borne pathogens (TBP) mirrors tick biting activity [3–6]. To acquire a host, *I. ricinus* employs a sit-and-wait strategy known as ‘questing’; the ticks climb vegetation and use sensory organs on the extended foretibiae to detect chemical cues from passing hosts [7, 8]. Questing activity is strongly determined by a range of climatic and microclimatic factors, particularly temperature and humidity (e.g. [9]) and, as a result, is affected by vegetation cover [10, 11] and interacting effects of landscape and host abundance (as reviewed by [12]). Hence, seasonal patterns of activity can vary widely from year to year. Longitudinal sampling, spanning months to years, has allowed the seasonal patterns of activity in different locations and between years to be well described [3, 4, 8, 9, 13].

In comparison to seasonal patterns, the diel activity of *I. ricinus* has not been well studied and no consensus has been achieved [14–16]. In laboratory experiments, negative phototaxis has been observed [7, 17], as well as darkness-induced mobility under a range of abiotic conditions [7, 18]. Furthermore, in the field in Richmond Park, UK, tick questing was negatively associated with light intensities [19], and a six-year study in Central Bohemia observed that tick questing began in spring when photoperiod exceeded an average of 12 hours, and ceased in autumn when photoperiod fell below 9 hours [20].

However, it is to be expected that climate and weather patterns, mediated by habitat, will also influence daily patterns of activity, but this has proved difficult to quantify. In Hungary, more ticks were observed in the three hours after sunrise compared to before sunrise, which led the authors to speculate whether light could initiate questing as there was no correlation with other environmental variables such as temperature [16]. Notably, the latter study’s plots were shaded by trees. In meadow habitat in Sweden, greater nocturnal activity was observed compared with daytime activity, but no discernible pattern was observed in nearby woodlands, and the availability of all stages was negatively correlated with temperature [14]. This habitat-specific behaviour was attributed to high mid-day saturation deficits in the more exposed meadow, exposing ticks to desiccation. In contrast, in open woodlands in the Netherlands, there was no significant difference in the number of larvae or adults questing in the day vs night, but significantly more nymphs were collected at night [21], and more dominant nocturnal activity was observed in plots with less vegetation coverage, implying a more exposed microclimate [21]. Furthermore, experimental observations support a nocturnal pattern of activity, driven by saturation deficits [22, 23]. On hill pasture in the United Kingdom, *I. ricinus* adults and nymphs were observed to be most active in the day and descended the vegetation at night [8, 24]. This diel behaviour was attributed to temperatures dropping below 10 °C at night, hence ticks descending into leaflitter. In contrast, another study recovered fewer ticks at mid-day in the UK in May, and this was associated with higher temperatures [23].

In other arthropod vector species, such as mosquitoes, where daily patterns of activity have been particularly well-studied, including factors driving and modulating biting behaviour (e.g. [25]), there is evidence that daily activity patterns influence pathogen transmission and the efficacy of control measures [26]. Variable patterns of tick questing activity therefore have potentially important epidemiological consequences. A better understanding of ecological drivers of the seasonal and daily activity patterns of *I. ricinus* is necessary to manage risk of tick bites and disease transmission to humans and other mammalian hosts. A clearer understanding of these behaviours and how they relate to climate may allow for mitigation strategies to be put in place, such as seasonal veterinary prophylactic treatments or behaviour modification to minimise exposure during periods of peak hazard. The aim of this study, therefore, was to observe the host-seeking activity of ticks throughout day (24 hours) and year, and relate this to different environmental parameters.

## 2 Methods

### 2.1 Study site

The study site was a 0.04km2 sampling zone within a 4-hectare section of unfenced deer sanctuary at Lyme Park (53.34, -2.04 (decimal degrees); Figure 1), a site located 15 km south-east of Manchester, England. The site is on a north-west facing slope, with an elevation of 250 – 270 m. The habitat was largely open lowland fen with underlying geology consisting of sandstone with coal measures and soils with low permeability. There is a westerly prevailing wind and an average annual rainfall of 600 mm to 700 mm. The deer sanctuary is home to approximately 150 red deer (*Cervus elaphus*), which are free to roam around the 550-hectare park and interact with the park’s resident populations of highland cattle (*Bos taurus*) and sheep (*Ovis aries*). Within the deer sanctuary there are also rabbits (*Oryctolagus cuniculus*), badgers (*Meles meles*) and a variety of small rodents. As a result, the vegetation of the survey site largely comprised typical fen species including sedges, rushes and grasses, with some “mown” areas of more intense deer activity.

**Fig. 1.**
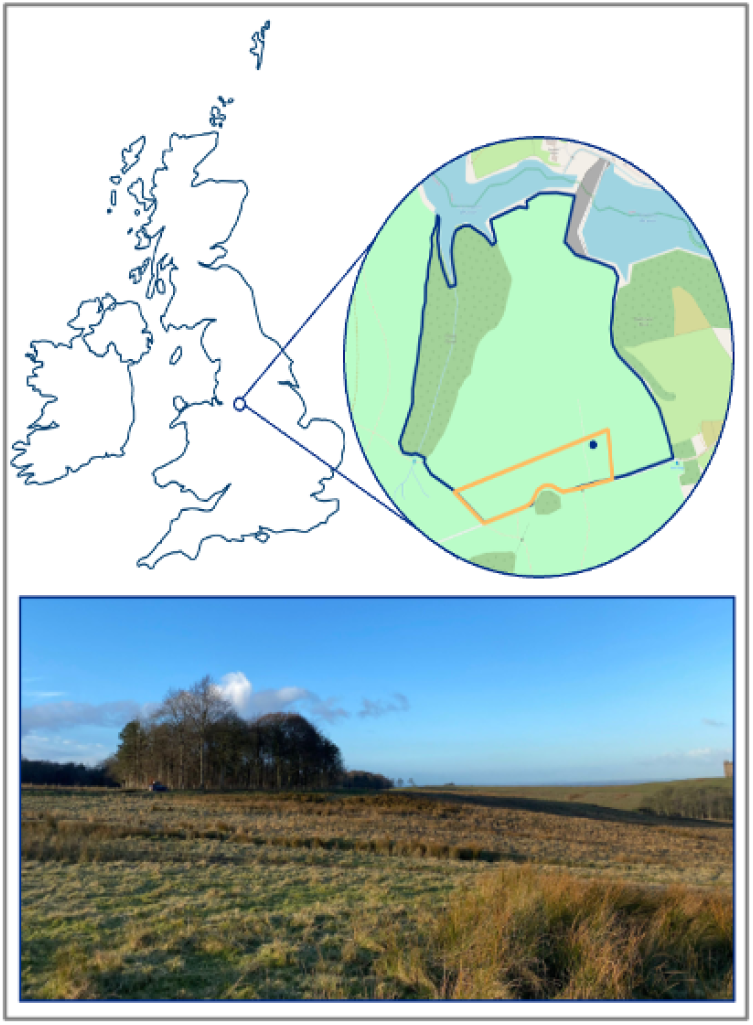
The location of the study site in the United Kingdom, with the deer sanctuary at Lyme Park outlined in blue, and the 0.04km2 sampling zone is in orange (OpenStreetMap, 2024). The bottom image represents the habitat of the study region, the photograph was taken from the blue dot presented in the study region map.

### 2.2 Assessment of diel and seasonal tick questing behaviour

Initial scoping surveys of the site, conducted from June to November 2021, indicated a high tick abundance. A subset of 300 ticks from these surveys were identified morphologically [27] as *I. ricinus*. In the main study conducted in 2022, to quantify daily and seasonal questing behaviour of ticks, blanket dragging was conducted over a 24-hour cycle, once a month. During each 24-hour cycle, the site was sampled every four hours at 08:00, 12:00, 16:00, 20:00, 00:00 and 04:00h (GMT). During darkness, head torches were used. At each of these 6 sampling periods within each 24-hour cycle, ten 10 m long blanket drags (transects) were conducted using a 1 x 1 m square of lightweight white cotton cloth pulled slowly over the ground [28] [29]. Before each 24-hour sampling cycle, the location and orientation of each transect was randomly selected within a 0.04 km^2^ subsection of the deer sanctuary using R statistical software (v.4.2.1; [30]). Each transect began at the randomised location and in the randomised orientation, except where there were obstacles, such as resting deer, whereby the closest safe path was chosen. At 5 m and 10 m of each transect, both sides of the cloth were inspected; adults and nymphs were removed using forceps and placed into an Eppendorf tube. A visual estimate of the number of larvae was made before they were removed and placed back into the environment. We state that the number of larvae is an estimate only due to their small size, that they tend to be recoverd in high-density clusters, and the speed at which they move, which in combination, make precise counts impossible under field conditions. Hereafter, all larvae quantities should be interpreted as estimates. All Eppendorf tubes containing ticks were kept chilled using frozen blocks until being returned to the laboratory within 24 hours of collectionto verify the number of nymphs and adults collected per transect. On some occasions, due to high winds and/or dark conditions, ticks escaped before being collected and these ticks were accounted for in the final numbers by noting these occurrences in the field as they occurred. A fresh blanket was used when the cloth was damp or dirty, or when a cluster of larvae was detected, to ensure no larvae were left on the cloth and that accurate recordings were taken at the next transect.

### 2.3 Environmental variables

To relate tick questing activity to environmental covariables, at the start of each transect, the atmospheric temperature and relative humidity (RH) 10 cm above ground level were recorded (Tinytag View 2 - TV-4500: Temperature ± 0.02 °C and ± 0.1% RH; Gemini Data Loggers, UK). The soil temperature was also recorded, by placing the thermistor probe 3 cm horizontally beneath the surface (Tinytag View 2 - TV- 4020 with a Thermistor Probe - PB-5001-1M5; Temperature ± 0.02 °C; Gemini Data Loggers, UK). However, no soil temperature data could be collected for two transects in August and October, respectively, because the thermistor probe broke. Data for these transects were excluded from statistical analyses.

Illuminance was recorded 50 cm above ground level (BT-881D Digital Illuminance Light Meter: Lux ± 4% *<*1,000 lux, ± 5% *<*1,000 lux, BTMETER, UK). All artificial light sources were covered or switched off when illuminance was recorded. For sampling sessions conducted in complete darkness, with no interruption of sunrise or sunset, the lux was recorded as *<*1 lux. Soil moisture was taken at three points along the transect (0, 5, 10 m), using a soil moisture probe (SM150T Soil Moisture Sensor and HH150 Moisture Meter: volumetric water content ± 3%; Delta-T Devices, UK). When readings were “Out of Range” due to high water content, the upper limit of 1,600 mV was assigned. In January and November, soil moisture was measured by dry weight analysis of three samples of soil from each transect taken at the same depth of the moisture meter. Gravimetric dry weight was then calibrated with the Moisture Meter in December 2022. However, visualisation of the soil data suggested that there was substantial variance and no meaningful relationship between measures taken using the soil moisture probe and estimates made from soil samples. Thus, the soil moisture estimates for January and November were excluded from further analyses. The vegetation height at three points of the transect (0, 5, 10 m) was also recorded using a 1 m metal ruler. An additional categorical system was employed to document the moisture on the vegetation; dry, light dew, heavy dew, light frost, heavy frost, wet (after rain). As this was a subjective variable, for analysis this was converted to a binary variable of dry or wet vegetation, where wet represented any form of water droplet on the vegetation. No frosts were observed.

In addition, temperature, precipitation, wind speed, wind gusts and cloud cover were collected for each of the sampling intervals (08:00, 12:00, 16:00, 20:00, 00:00, and 04:00) at the coordinates (53.34, -2.04 (decimal degrees)). from the Visual Crossing Corporation[31].

Saturation deficit (SD) was calculated using the temperature (T) and relative humidity (RH) at 10 cm [22]. To allow a comparison of ticks questing in the day or at night, the final illuminance recordings were converted to binary measurements, where records equal to or above one lux were defined as light, and those below this threshold were defined as dark. The threshold of one lux was chosen to account for urban light pollution from nearby cities [32], the presence of a full moon [33] and the uncertainty in measurements. Continuous measures based on the raw illuminance data were not used in analysis as these can vary minute-by-minute, e.g., due to cloud cover.

### 2.4 Statistical analyses

Data were deposited in the VecDyn database [34] (https://vectorbyte.crc.nd.edu/vecdyn-detail/951) and accessed using the ohvbd R package [35] for analysis. The full dataset and R code required to replicate these analyses are also available in an OSF repository (https://osf.io/hymwb/overview?viewonly=5344e8b19918479882d1a073c2c6c3b1) NOTE TO COPYEDITOR - this link is blinded for review and must be replaced with a public link before publication.

The seasonal patterns of activity of adults, nymphs and larvae were plotted and then assessed using cardidates::peakwindow R function (v.0.4.9; [36]) on the sum of each instar to identify any peaks in the time series data.

The influence of environmental factors on the density of questing nymphs was tested using generalised linear mixed models (GLMMs); the density of questing nymphs is the most epidemiologically important instar stage and nymphs were observed in sufficient numbers for analysis. As is typical with ecological data, the response variable comprised count data that were overdispersed (variance:mean = 9.24) but were not zero-inflated (32.8% zeroes). Therefore, competing negative binomial and log-normal Poisson models with and without month as a random effect (lme4::glmer.nb v.1.1-28; [37]), and a mean-variance model incorporating month as both a random effect and a dispersion factor (glmmTMB package; [38]) were compared by Akaike Information Criterion (AIC) and residual diagnostics (DHARMa and performance packages; [39], [40]). The mean-variance model with sampling month as both a random effect and dispersion factor, and an observation-level random effect, outper-formed all others based on AIC (mean-variance model AIC = 2620, compared with AIC in the range of 2631-3095 for all other models) and residual diagnostics. Including month as a dispersion factor allows dispersion to vary by month, while including month as a random effect allows the mean to vary with each month.

To avoid collinearity effects, correlations between the continuous explanatory variables were checked using a Spearman’s correlation matrix (Hmisc::rcorr v.4.7-2; [41]). If two variables had over 70% correlation, only one was taken forward. Final models were also assessed for collinearity, and a variance inflation factor (VIF) below 5 was considered acceptable (car::vif v.3.1-0; [42]). Explanatory variables were log-transformed and z-scaled, with the exception of light levels (categorical light vs. dark), for model input.

A multivariable model containing the non-correlated explanatory variables was constructed and residuals checked once more. The model was finally compared with a null model containing only the random effect factor in a likelihood ratio test (stats::anova v.4.1.2 (test = “chisq”); [30]). The value of the random effects was assessed by comparing the final model with and without random effects, using likelihood-ratio tests.

Vegetation moisture and soil moisture were excluded from analyses due to missing data for January and November. The month of June was exclude from all analyses, because no sampling took place due to COVID-19 restrictions. above.

## 3 Results

### 3.1 Seasonal patterns of activity differed between instars

Ticks were collected every month, except in June 2022. Over 8,000 ticks were counted (Figure 2, Table 1, Supplementary Figure S1); larvae were most abundant (approx. 5,034; 62.84%), followed by nymphs (2,715; 33.89%) and then adults (262; 3.27%). The three instars had different seasonal patterns of activity. Peak window analysis identified one peak in the density of questing larvae in July with a median of 10.5 larvae / 10 m^2^ (range 0-approx. 258), implying a unimodal pattern of activity (Figure 2). Two peaks were identified for nymphs; a small spring peak in April, with a median of 2 nymphs / 10 m^2^ (range 0-16), then a larger peak in November, with a median of 10 nymphs / 10 m^2^ (range 0-37)(Figure 2). A bimodal pattern of activity was also observed for adults, with a small spring peak in March with a median of 0 nymphs / 10 m^2^ (range 0-2) and a larger peak in September with a median of 10 nymphs / 10 m^2^ (range 0-6) (Figure 2). However, note that the low number of adults recovered with each transect inflates uncertainty in the timing and magnitude of these peaks. At least one instar stage was collected each month, with only nymphs being collected in January and only larvae in July (Figure 2, Table 1).

**Fig 2.**
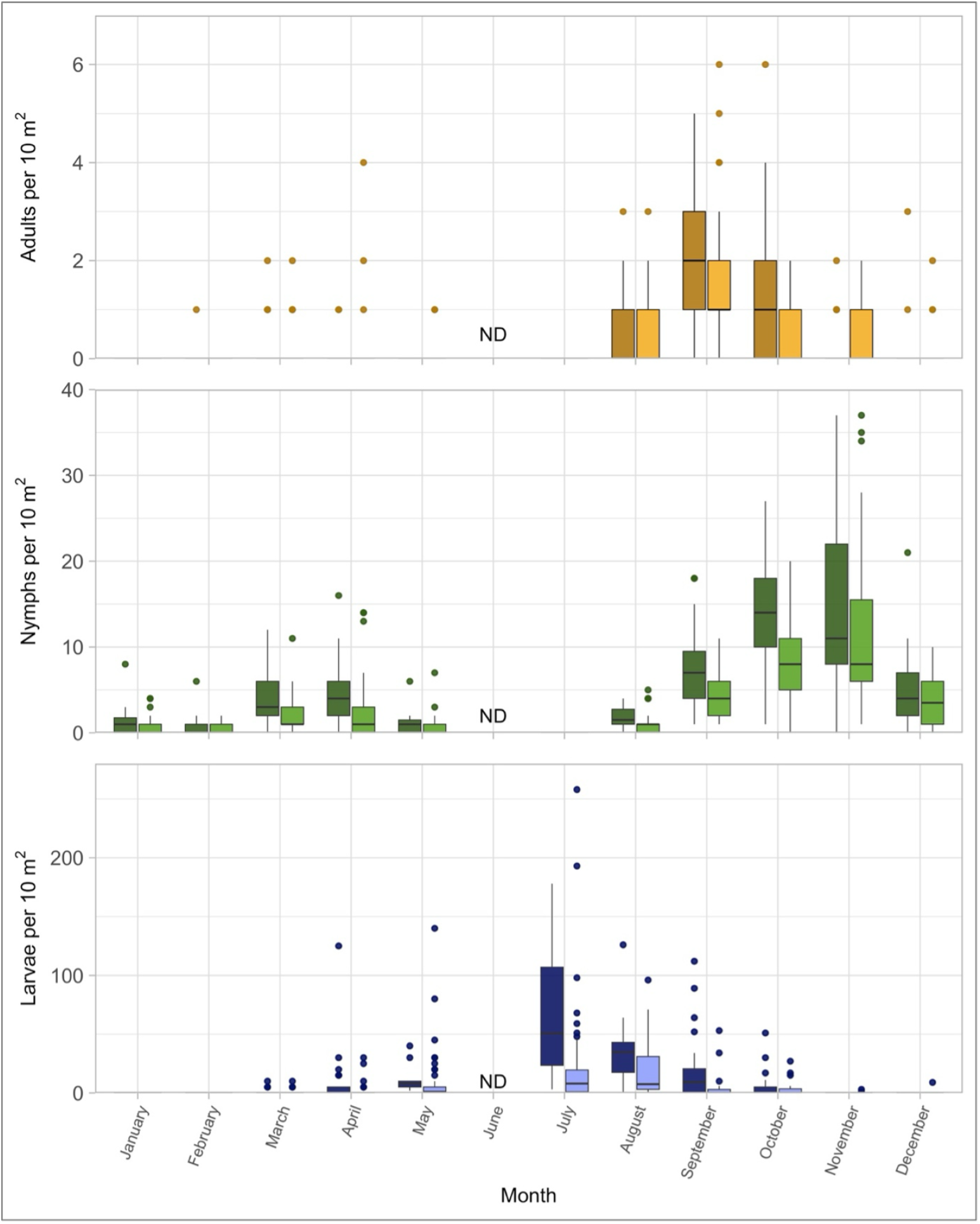
The density of questing of adults, nymphs and larvae, counted in light and dark conditions ach sampling month. The darker colour (left bar) represents ticks counted where light levels under one lux, and the lighter colour (right hand bar) represents light levels over one lux. Boxes esent the interquartile range of the data, with a line at the median value. Whiskers extend to minimum and maximum values, with outliers (points more than 1.5 times the interquartile range e the third quartile (upper limit of the box) or below the first quartile (lower limit of the box)) esented by filled circles.

**Table 1.** The total number of larvae, nymphs and adults collected within the 24-hour sampling session each month. For larvae, these approximate total numbers due to their small size.

| Month | Start Date | Larvae | Nymphs | Adults |
| --- | --- | --- | --- | --- |
| January | 13-01-2022 | 0 | 56 | 0 |
| February | 11-02-2022 | 0 | 28 | 1 |
| March | 21-03-2022 | 40 | 191 | 13 |
| April | 10-04-2022 | 310 | 200 | 9 |
| May | 21-05-2022 | 597 | 47 | 3 |
| July | 18-07-2022 | 1,823 | 0 | 0 |
| August | 08-08-2022 | 1,362 | 72 | 32 |
| September | 13-09-2022 | 649 | 357 | 116 |
| October | 03-10-2022 | 240 | 672 | 60 |
| November | 12-11-2022 | 4 | 837 | 20 |
| December | 03-12-2022 | 9 | 255 | 8 |

### 3.2 Diel activity – questing tick density was highest at night

Ticks of all three instar stages were collected at all sampling times (08:00, 12:00, 16:00, 20:00, 00:00 and 04:00). However, the daily activity of adult, nymph, and larval ticks varied between months (Supplementary Figures S2 - S4). Fewer adult ticks were collected at 08:00, 12:00 and 16:00 h sampling times, compared to 20:00, 00:00 and 04:00 h sampling times, from February to October (Supplementary Figure S2). Similar results were found for nymphs (Supplementary Figure S3). The number of larvae collected varied across the year. From April to October, more larvae were collected at 20:00, 00:00 and 04:00 h sampling times, but this pattern of activity was not observed in the remaining months (Supplementary Figure S4), with the highest collection (n = 258) occurring at 20:00 in July.

#### 3.2.1 Environmental factors – questing activity was observed under a wide range of conditions

All instar stages quested across a range of environmental conditions, between 1.6 and 40.3 °C, and between 24% and 100% RH (Figure 3). Nymphs were the only instar stage found in January, when the mean temperature across the 24 hour sampling period was 3.51°C [31]. Adults were first collected in February, and larvae in March when mean temperatures were 3.91 °C and 7.28 °C respectively [31]. The nymphs were found at the lowest transect temperatures (1.6 °C in February at 20:00), followed by adults (2.1 °C in February at 20:00), then larvae (3.3 °C in April at 04:00). Larvae, however, were the only instar stage found questing in July, when transect air temperatures were as high as 40.3 °C, and at relative humidities as low as 24% (Figure 3). No adults or nymphs were found questing when the mean daily temperature exceeded 25 °C. The blanket drags with the highest adult and nymph yields (denoted by darker points in Figure 3) were between 7 and 20 °C with humidities greater than 70% RH.

**Fig. 3.**
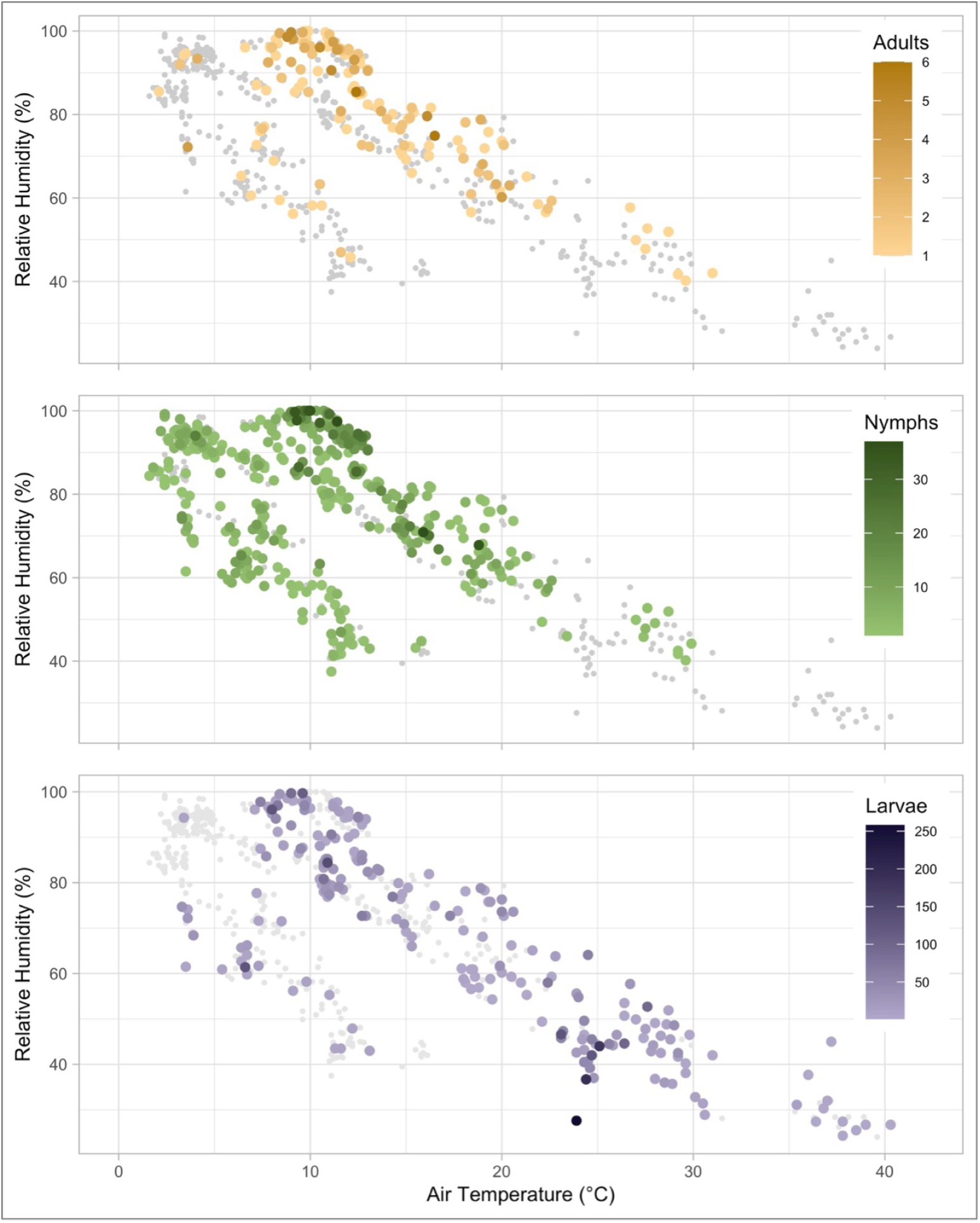
The density of questing adults, nymphs and larval ticks collected per 10 m^2^ of tick dragging under different environmental conditions. The temperature and relative humidity were recorded at 10 cm above ground level. Coloured circles represent the different life stages of ticks collected, with darker colours reflecting greater density of ticks, and smaller grey circles representing the sampled conditions where that life stage was absent.

#### 3.2.2 Environmental factors – questing nymph density is associated with saturation deficit and light levels

The multivariable GLMM, included variables with correlation coefficients below 0.7 (saturation deficit, light level, soil temperature, vegetation height, and wind speed) as fixed effects, and month as a random effect (Table 2). The model predicted that density of questing nymphs was around 30% greater under dark conditions compared to light (IRR = 1.3), and decreased by around 20% for each standard deviation decrease in log(saturation deficit+1) (IRR = 0.8; Table 2; Figure 3; Supplementary Figures S2 - S5). The same broad observation was made (but not explicitly modelled) for larvae, and this night-time activity appeared to be most dominant in July when an estimated median of 50.5 larvae/ 10 m^2^ was found at night versus 8 larvae / 10 m^2^ in the day. No strong daily pattern of activity in relation to light and dark was observed for adults, potentially due to the low numbers recovered.

**Table 2.** GLMM multivariable analysis of the relationship between different environmental factors and the density of questing nymphs in Lyme Park, UK, in 2022. Month and observation-level (OLRE) were included as random effects, with a dispersion formula by month (negative binomial, log link). Incidence risk ratio (IRR = exp(coefficient estimate)) and 95% confidence interval (CI), model coefficient estimate and standard error (SE), z-score and p-value are presented. Significant results are highlighted in bold.

| Predictor | IRR (95% CI) | Estimate (SE) | Z | P |
| --- | --- | --- | --- | --- |
| Intercept | - | 0.775 (0.341) | 2.28 | **0.023** |
| Saturation deficit (log+1 scaled mmHg) | 0.788 (0.664-0.935) | -0.239 (0.088) | -2.73 | **0.006** |
| Light level (Dark) | 1.296 (1.063-1.580) | 0.259 (0.101) | 2.56 | **0.0105** |
| Soil temperature (log+1 scaled $^{\circ}C$) | 1.106 (0.860-1.424) | 0.101 (0.129) | 0.79 | 0.432 |
| Wind speed (log+1 scaled mph) | 1.011 (0.902-1.134) | 0.011 (0.059) | 0.19 | 0.849 |
| Vegetation height (log+1 scaled cm) | 1.054 (0.967-1.150) | 0.053 (0.044) | 1.20 | 0.231 |

Dispersion estimates did not vary significantly by month, except September (1.63 (0.85 SE), z = 1.93, p = 0.054) and October (1.84 (0.93 SE), z = 1.97, p = 0.049) where dispersion was slightly higher relative to a January baseline.

The final model outperformed a null model (*χ*^2^(6, N=658) = 45.4, p = *<*0.001), and a model with only fixed effects (*χ*^2^(2, N=658) = 196, p = *<*0.001). Residuals were uniform, with no dispersion issues, outliers, nor zero-inflation issues. There was minor heteroscedasticity in residuals vs fitted values and minor deviations from uniformity in the residuals vs factors. There was no autocorrelation with sampling period although some weak seasonality was evident in the autocorrelation function on the monthly data. Introduction of the dispersion formula removed the majority of the heteroscedasticity, but a positive trend in the residuals vs fitted values remained, suggesting that the model tended to underpredict at lower fitted values, and overpredict at higher fitted values.

## 4 Discussion

This study quantified the daily activity of *I. ricinus* over complete 24-hour periods and throughout the year at a site in Northern England – whereas previous studies omitted night or winter observations [14, 16]. A daily rhythm of nocturnal activity was observed consistently, whereby the density of questing nymphs was predicted to be 30% greater at night, with some variation throughout the year. This is consistent with observations of ticks in open habitats in other European regions [14, 21] and a reduction in questing activity at mid-day has been observed elsewhere under both field [19, 23] and experimental conditions [22]. Such daily rhythms have also been documented in other parasites and vectors, and can arise due to endogenous or exogenous factors (reviewed by [43]). The study presented here sought to identify the environmental factors driving seasonal and diel patterns of activity in ticks.

There are some apparent inconsistencies between studies observing the daily behaviour of ticks, but these may be explained by other environmental covariables influencing microclimate. For example, the daily pattern of activity has been found to vary between life-cycle stages [14, 21] over seasons [16, 44], and between locations and habitats [14]. This highlights a potential limitation of most studies, including our own, which typically have a restricted geographic scale of sampling, incorporating a relatively homogenous habitat. Although this has the benefit of potentially reducing environmental confounders, the environmental conditions captured in the dataset may be less variable than more heterogeneous sites, or future multi-site studies. Mapping the parameter space encountered during sampling, as we have done here, may facilitate between-study comparisons. However, the variability of outcomes between studies nevertheless highlights their individual limitations in capturing the complexity of the study system and the need for a more comprehensive mechanistic understanding of tick ecology.

Despite between-study variability, the broad interpretations of previous observations of ticks throughout Europe are consistent with the current analyses. Diel patterns of nymph activity were negatively, and significantly, associated with both light levels and saturation deficit. Similar trends towards nocturnal activity were also qualitatively apparent for larvae, while data for adults were too sparse to draw inference. Ixodid tick mortality is highly dependent on water loss, which is regulated by the atmospheric saturation deficit [45]. At night, the cooler temperatures and higher relative humidities, hence lower saturation deficits, may provide less desiccating conditions more frequently for questing [16, 21, 22]. This moderating effect of saturation deficit is evident in our observed seasonal patterns of tick activity. There was no obvious seasonal shift from an exclusively diurnal to nocturnal pattern of activity here, but in the months where there was little variation in the saturation deficit throughout the day, such as January, February and December, the nocturnal pattern of activity was less evident. In contrast, in months where midday saturation deficits were high, such as July, there was a dominant pattern of nocturnal activity.

Olfactory and kinetic cues caused by a high density of nocturnal rodent hosts have been suggested as drivers for tick behaviour [7, 8], and there are diverse examples of other parasites whose activity patterns coincide with host availability, thus maximising transmission opportunities (reviewed by [43]). At Lyme Park, small rodents and deer were present, but not recorded and hence a correlation between host density and activity and tick questing cannot be established. However, the nocturnal patterns of activity observed in ticks, corresponds to nocturnal patterns in small mammals in the UK [46] and the nocturnality observed in deer in areas of high human activity [47, 48]. Consequently, both climatic conditions and host activity may influence the daily behavioural patterns of ticks.

The results of the present study suggest that there is greater risk of tick exposure in warm weather (nymph densities tended to be highest in the warmer spring and autumn months), at night and around dawn, and when saturation deficits are lower. This could have important epidemiological considerations. For example, in the summer there are increased evening recreational activities such as camping, which would increase human exposure. Similarly, it is advised that in warm weather, pets should be exercised during the cooler parts of the day, including early morning and late in the evening, which could coincide with higher densities of questing ticks[23].

The seasonal pattern of tick activity observed in Lyme Park resembles patterns previously recorded for *I. ricinus* in the UK [3, 4, 49, 50]. The relatively high numbers of questing nymphs and adults observed in spring is likely to be due to the emergence of ticks from behavioural quiescence as temperatures rise [3, 51, 52], while the autumnal peak in activity is likely to represent attempted feeding of the newly moulted cohort of ticks. The often low autumn peak seen in the UK has been attributed to cooler summer temperatures delaying development [3]. However here, spring-fed juvenile ticks may have fed and moulted into their next instar stage by autumn increasing the observed density of questing ticks between September and November. The low numbers of questing ticks commonly reported in mid-summer is likely to be driven to the environmental conditions; if the saturation deficit rises above an estimated 4.4 mmHg, *I. ricinus* exhibits positive geotropism behaviours to avoid desiccation [3, 9, 17]. Other longitudinal studies have observed this bimodal pattern of tick activity in regions, altitudes and years where there is high moisture stress during the mid-summer period [9, 52–54].

Although the larval pattern of activity followed the typical distribution observed throughout the UK, the timing of the peak activity was unexpected as it coincided with high saturation deficits exceeding 40 mmHg, which is well above the threshold considered suitable for larval questing [3, 9, 10, 17]. Our approximation of larvae numbers undoubtedly introduces additional uncertainty to the absolute quantities, however, the same methods were applied by the same researcher throughout the study and thus generate a useful longitudinal dataset of seasonal patterns of abundance. Larvae are thought to be most at risk of dessication due to their small size, restricting questing activity to periods of optimal environmental conditions (as reviewed by [55] and [56]). Our data challenge this generalisation, suggesting that questing behaviour is driven by interacting factors, some of which were not measured during this study but which may act antagonistically, resulting in questing during suboptimal environmental conditions. For example, it is it possible that the behaviour was driven by starvation [57, 58] leading to questing under sub-optimal conditions. We hypothesise that larvae questing under conditions of high saturation deficit would have lower lipid reserves than those questing under less desiccating conditions. Lipid analysis on a larger sample of ticks would be required to quantify these differences, if present. Gray [59] also proposed alternative hypotheses for a unimodal pattern of larval activity, including more rapid “activation” of ticks in open habitat, and a late spring peak resulting in a smaller autumn peak.

Beyond challenging our assumptions on the lower temperature thresholds for tick activity (ticks were observed questing at 1.6 *^◦^*C in this study, and at sub-zero temperatures in initial scoping surveys at the same site in 2021; unpublished data, M. Noll), the observations reported here have important implications for tick ecology research, since biases introduced by the timing and conditions of tick sampling may be underappreciated. The density of questing ticks was lower at daytime sampling times typically represented in tick surveys (08:00, 12:00 and 16:00) compared to atypical sampling times (20:00, 00:00 and 04:00); up to eight times more larvae were collected outside of typical sampling hours. It should therefore be noted that the increase in the recorded density of questing ticks at night is substantial, considering that one could expect more type two errors at night (false negatives) due to the small size of ticks, especially larvae (1 – 2 mm), rendering them difficult to see under artificial light [44]. Therefore, it is likely that current blanket dragging practices (sampling during daylight only) severely underestimate the active tick population. Furthermore, there was a higher density of questing nymphs when the vegetation retained water droplets, which were often caused by dew. The presence of dew on vegetation may increase the relative humidity and improve the microclimate to favour questing [10]. This was an unusual finding, as tick sampling is seldom performed during or after rain due to wet vegetation [16, 19]. Therefore, the current standard sampling protocols may need to be adjusted to encompass sampling with wet vegetation (both dew and rain induced), to extend our understanding of tick behaviour under these conditions. To mitigate the potential impacts of wet conditions on sampling efficiency, researchers could regularly replace damp blankets with dry blankets, as was done in our study. Given that sampling throughout the night has practical, logistical and safety limitations, simply including early morning and late evening sampling would help to increase the accuracy of abundance estimates [23].

Due to the dependency of tick development and activity on climatic factors, it is likely that there will be seasonal changes in the patterns of tick activity with climate change [60]. Any increase in the synchrony of larval and nymphal questing activity is of concern due to co-feeding between the instars contributing to direct pathogen transmission [61, 62]. If nocturnal questing is driven by the avoidance of desiccating midday conditions, then nocturnal patterns of activity may become more extreme, further increasing the opportunity for co-feeding between uninfected larvae and infected nymphs. However, if such changes in environmental conditions are at odds with circadian rhythms (which are, as yet, unexplored in ticks), then non-linear patterns of change in activity may arise, such as bimodal diel activity.

Further work is required to investigate the environmental and chronobiological drivers of host-seeking behaviour in ticks, and a recently-published guide signposts researchers new to this expanding field of research [63]. Our study highlights a number of limitations of such research, including the narrow geographic range, challenges of sampling with a high temporal resolution and minimising sampling error, and narrow range of species observed. Where possible, experiments and observations should be repeated at multiple sites, or using ticks sourced from multiple sites, since there is evidence for intraspecific variation of tick questing behaviour based on climate of origin [64], and include a range of species, since the rate of water loss is species-specific, possibly due to differences in cuticle permeability and spiracle morphology influencing transpiration rates [65]. Additionally, nutrition has been shown to influence host-seeking behaviour in mosquitoes [25], thus further work should also strive to incorporate nutrition status, where possible. For example, assessing the lipid content and age of the tick when collected, may provide insight into how starvation influences behaviour, but requires a large sample size. Finally, a negative influence of pathogens on tick fitness (if present) has been predicted, in a modelling study, to be a key parameter driving tick-borne pathogen transmission dynamics [66]. Thus, investigating the infection status of the tick would provide greater insight into how this influences their environmental tolerance [67–69].

Our empirical model identified two key drivers of the density of questing nymphs in an open habitat, but tended to slightly overestimate nymph density at high fitted values, and underestimate nymph density at low fitted values. We hypothesis that this may be related to seasonal and inter-annual tick population dynamics moderating the density of questing ticks. Where estimates of tick hazard are required for dynamic public/veterinary health risk assessments, mathematical models explicitly tracking tick population dynamics and behavioural response to environmental conditions may offer greater resolution and certainty (e.g. [70]).

## 5 Conclusion

The seasonal and daily activity of ticks observed in this study, driven by light levels and saturation deficit, has generated new insights into the drivers of tick behaviour and tick bite exposure risk, and provides a foundation for further work. Our results, interpreted in the context of previous work, evidence that both abiotic factors, such as the saturation deficit, and biotic factors, including host abundance, influence the daily and seasonal patterns of tick activity. Furthermore, our observations challenge recognised thermal limits of tick activity, indicating a potential risk of year-round exposure to ticks in temperate regions such as the UK, with elevated risk during the night and at dawn, and when daytime conditions result in high saturation deficit. This has implications for those managing tick hazards and potential exposure to tick bites.

## Supplementary information

Supplementary Figures are appended.

## Declarations

### Ethics approval and consent to participate

Ethical approval was granted by the University of Liverpool’s Veterinary Research Ethics Committee (Approval No.: VREC1099, granted: 08/06/2021). Informed prior consent was obtained from the land manager.

### Consent for publication

Not applicable

### Availability of data and materials

All data generated by this study have been deposited in the VecDyn database [34] for open use: https://vectorbyte.crc.nd.edu/vecdyn-detail/951 [Note that the dataset is under embargo until acceptance of this manuscript for publication].

### Competing interests

RW has previously received funding for research from MSD Animal Health UK.

### Funding

MN’s PhD studies were supported by MSD Animal Health UK. HRV was supported by the Biotechnology and Biological Sciences Research Council [BB/W016621/1] and by the Defra-UKRI One Health VBD Hub [BB/Y008766/1] during the analyses, data curation, and final preparation of this manuscript.

### Authors’ contributions

Conceptualisation; MN, HRV. Methodology; MN, RW, BLM, HRV. Formal analysis; MN, HRV. Investigation; MN, HRV. Resources; MN, HRV. Writing - original draft; MN, HRV. Writing - review & editing; MN, RW, BLM, HRV. Visualisation; MN. Supervision; RW, BLM, HRV. Project administration; BLM, HRV. Funding Acquisition; RW, HRV.

## Acknowledgements

We are grateful to Chris Dunkerley and the National Trust for permitting site access, to Sarah Kelly at the VBD Hub for data curation support, to Francis Windram at the VBD Hub for support with the ohvbd package, and to those who assisted with field data collection: Alice Coleman, Catherine Hewitt, Daisy-Grace Dunn, Isabella Endacott, Jordan Rolfe, Molly Brock, Rocio Checa-Herraiz, William Newland.

## Supplementary information

**Fig. S1.**
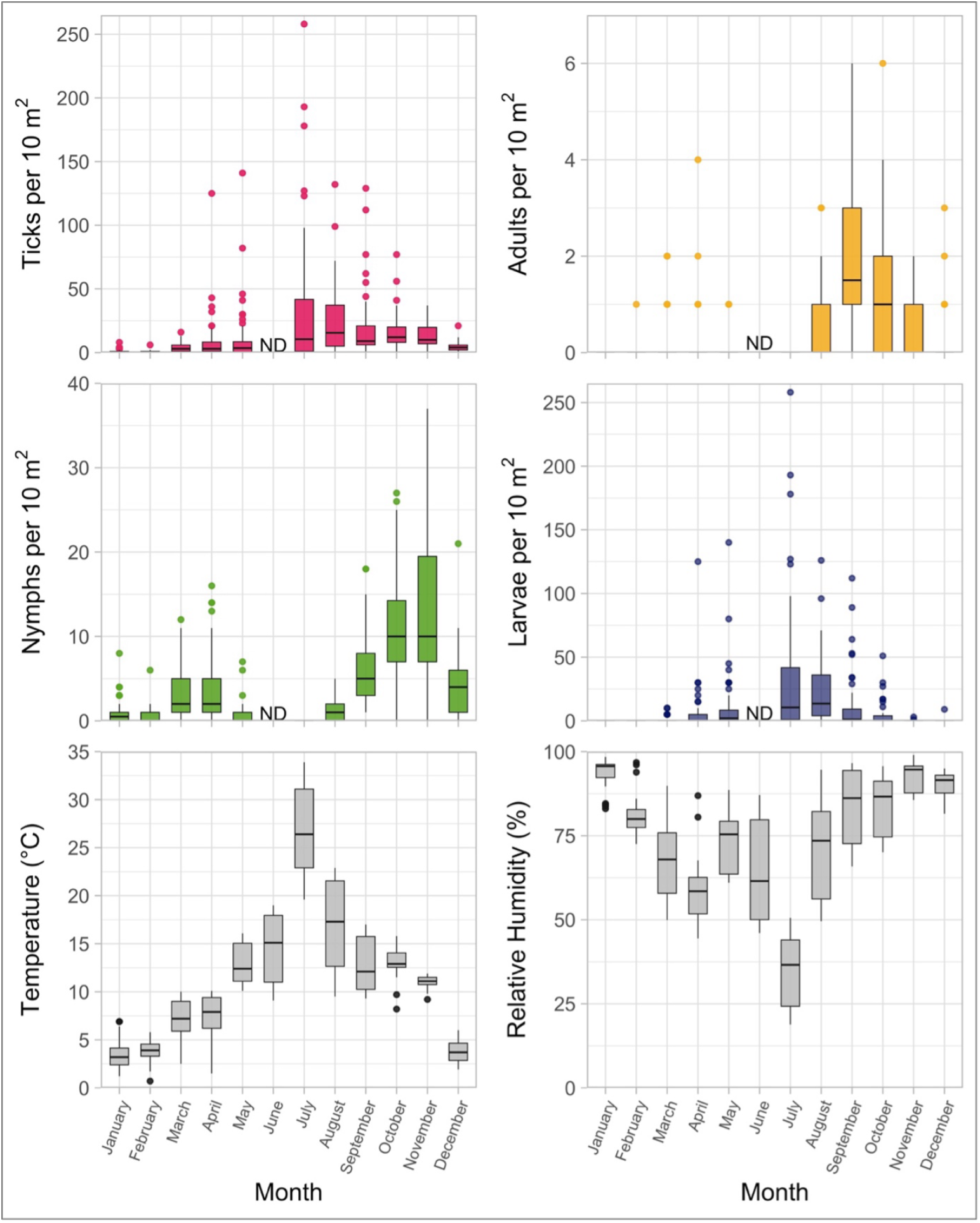
The density of questing ticks collected by blanket dragging at each transect in the 24-hour sampling cycles from January – December 2022 at Lyme Park, UK, and the air temperature and relative humidity during each sampling interval, taken from Visual Crossing (Visual Crossing Corporation, 2024). ND refers to no data, where no sampling took place in the month of June. Boxes represent the interquartile range of the data, with a line at the median density of ticks collected. Whiskers extend to the minimum and maximum values, with outliers represented by filled circles.

**Fig. S2.**
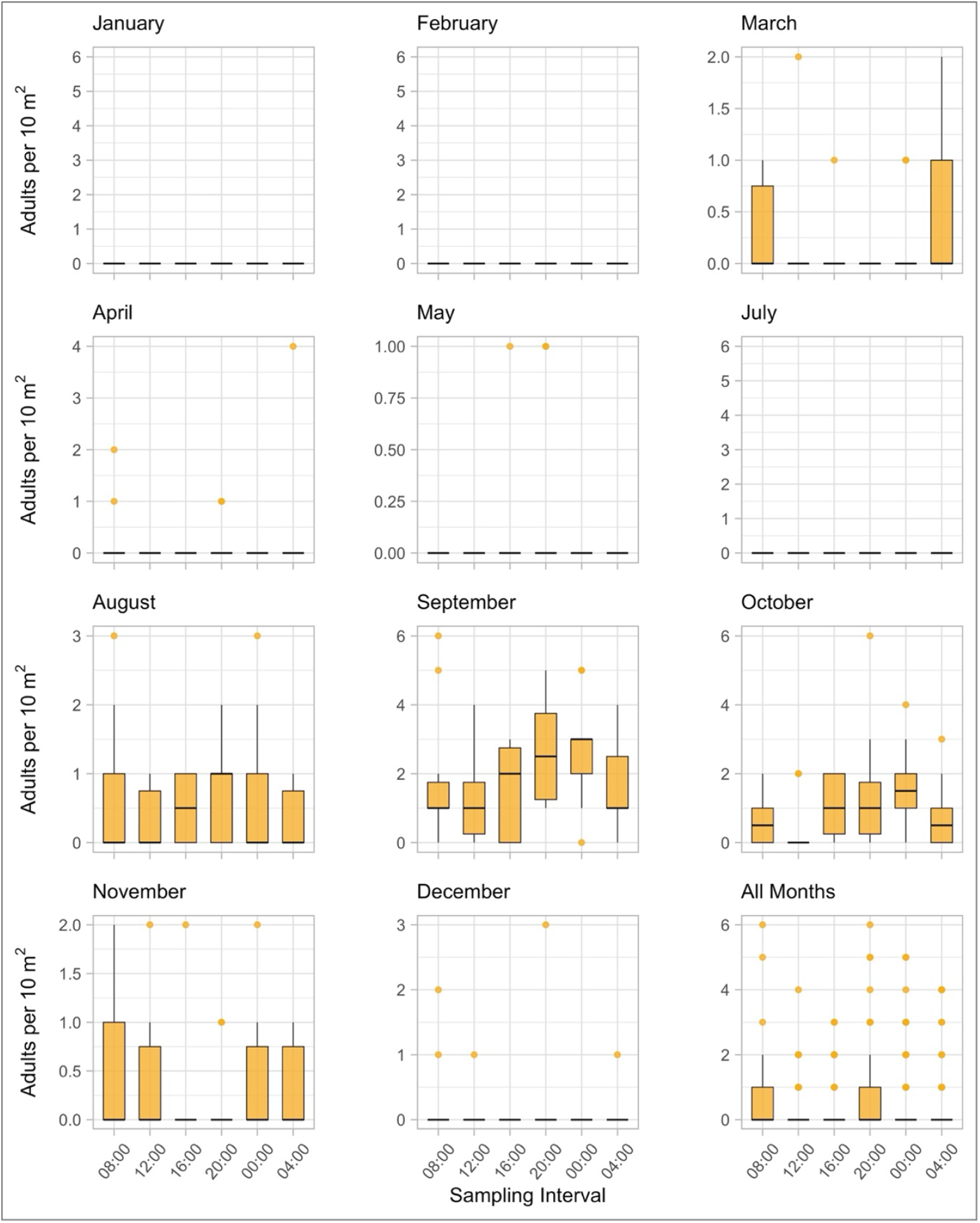
The density of questing adult ticks collected per 100 m^2^ of blanket dragging at each of the sampling periods [08:00, 12:00, 16:00, 20:00, 12:00, 04:00 (Greenwich Mean Time)] for individual sampling months, and a total of all months. Boxes represent the interquartile range of the data, with a line at the median value. Whiskers extend to the minimum and maximum values, with outliers represented by filled circles.

**Fig. S3.**
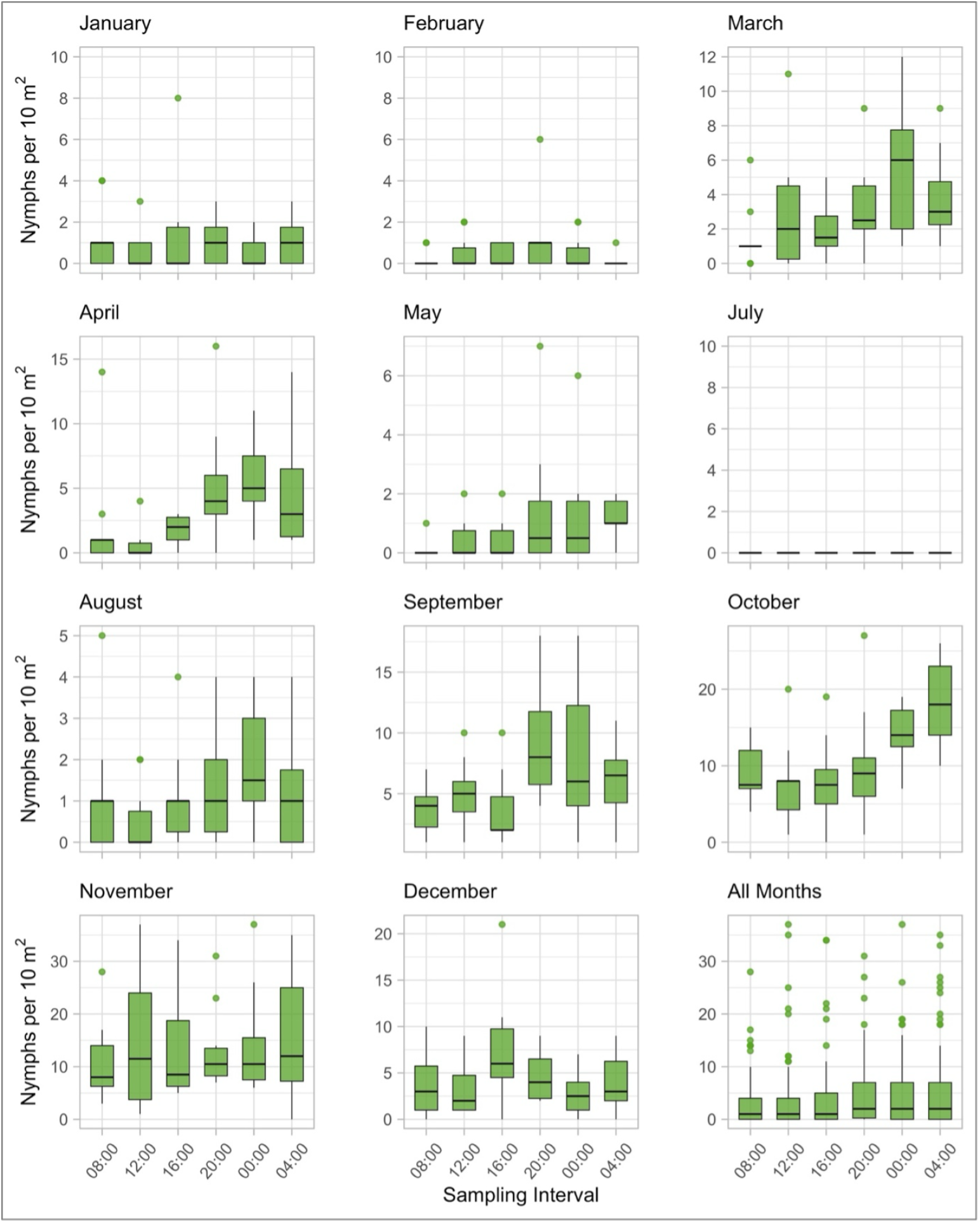
The density of questing nymphal ticks collected per 100 m^2^ of blanket dragging at each of the sampling periods [08:00, 12:00, 16:00, 20:00, 12:00, 04:00 (Greenwich Mean Time)] for individual sampling months, and a total of all months. Boxes represent the interquartile range of the data, with a line at the median value. Whiskers extend to the minimum and maximum values, with outliers represented by filled circles.

**Fig. S4.**
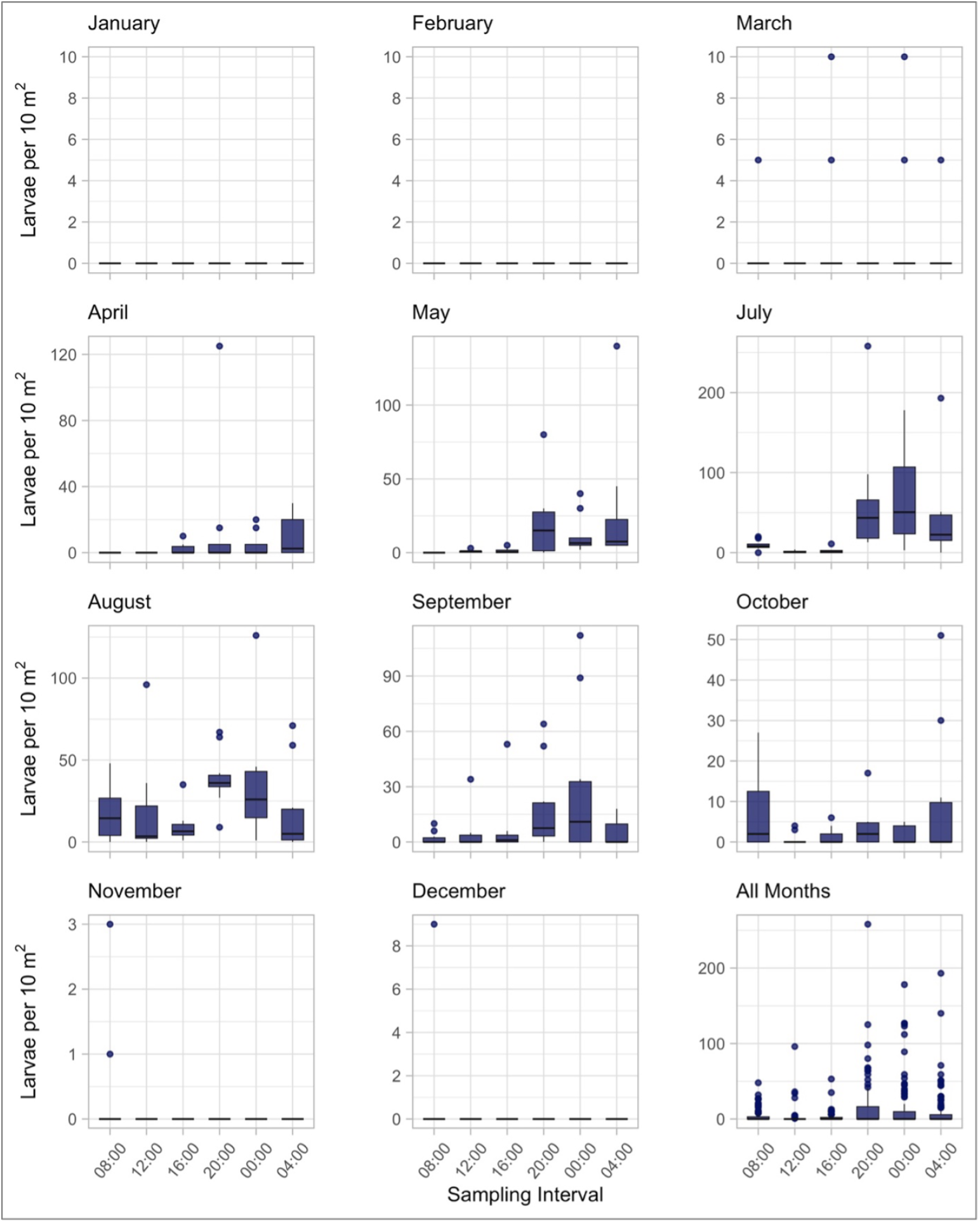
The density of questing larval ticks collected per 100 m^2^ of blanket dragging at each of the sampling periods [08:00, 12:00, 16:00, 20:00, 12:00, 04:00 (Greenwich Mean Time)] for individual sampling months, and a total of all months. Boxes represent the interquartile range of the data, with a line at the median value. Whiskers extend to the minimum and maximum values, with outliers represented by filled circles

**Fig. S5.**
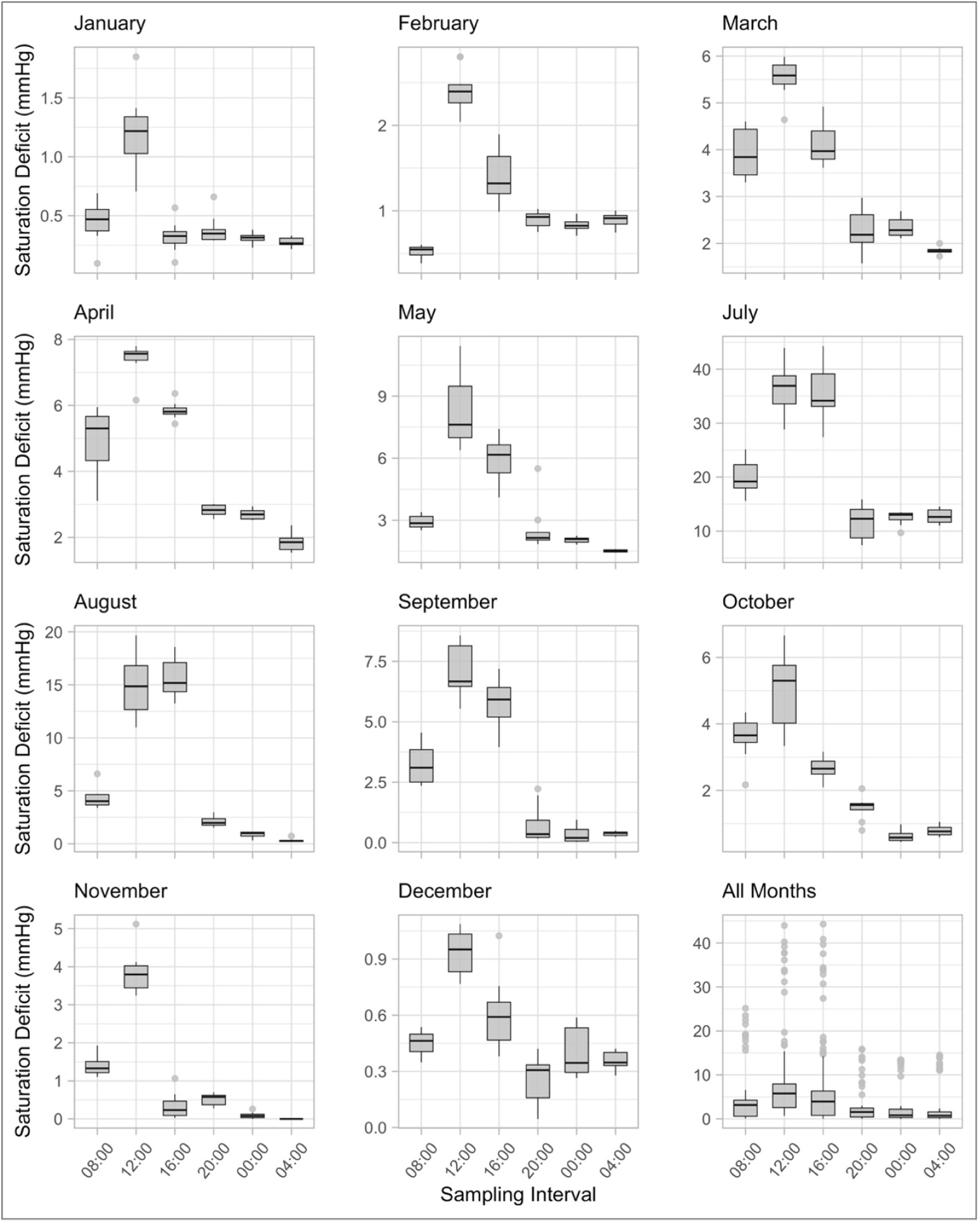
The saturation deficit (mmHg) at each transect in each of the sampling periods [08:00, 12:00, 16:00, 20:00, 12:00, 04:00 (Greenwich Mean Time)] for individual sampling months, and all months combined. Boxes represent the interquartile range of the data, with a line at the median value. Whiskers extend to the minimum and maximum values, with outliers represented by filled circles.

## Notes

### Competing Interest Statement

MN's PhD studies were supported by MSD Animal Health UK. RW has previously received funding from MSD Animal Health UK

### Summary of Updates

Restructuring of the manuscript as requested by reviewers. Analyses simplified to only include the full GLMM (backward stepwise selection of a reduced model removed). ohvbd package used to access data in the VecDyn repository.

